# EVOLUTIONARY TRENDS IN SPECIES COMPLEX DIAGNOSED BY CYTOGENETIC POLYMORPHISM: THE CASE OF *Hypostomus ancistroides* (SILURIFORMES, LORICARIIDAE)

**DOI:** 10.1101/2020.09.22.308601

**Authors:** Dinaíza Abadia Rocha-Reis, Karina de Oliveira Brandão, Jorge Abdala dos Santos Dergam, Rubens Pasa, Karine Frehner Kavalco

**Affiliations:** Laboratory of Ecological and Evolutionary Genetics - LaGEEvo, Institute of Biological and Health Sciences, Federal University of Viçosa, Campus Rio Paranaíba, Brazil; Department of Anatomy and Embryology, Leiden University Medical Center, Leiden, Netherlands; Laboratory of Molecular Systematic “Beagle”, Department of Animal Biology, Federal University of Viçosa - Campus Viçosa, Brazil

**Keywords:** Constitutive heterochromatin, Phylogeny, *Hypostomus*, Karyotypic evolution, Ribosomal sites

## Abstract

Hypostominae is a subfamily of Loricariidae with great variation in color characters and external morphology. The genus *Hypostomus* presents the largest number of species ever karyotyped, with *Hypostomus ancistroides* characterized as a group of cryptic species. In the 15 natural populations of *H. ancistroides* studied, there are 15 different karyomorphs, with variations in diploid number, sex chromosome systems, and markers, such as C-banding and location of ribosomal cistrons. The objective of this work was to present molecular and chromosomal data of four new populations of *Hypostomus ancistroides* and to discuss the observed evolutionary trends for this group of a cryptic complex of species. We analyzed specimens from four sampling points in the Tietê, Mogi-Guaçu, and Grande river basins, all in the state of São Paulo, southeastern Brazil. We performed techniques such as the detection of constitutive heterochromatin and ribosomal sites (5S and 18S), in addition to phylogenetic analyses. All specimens presented 2n=68 chromosomes without supernumerary elements or sex-related heteromorphisms. However, each population has a different karyotype with unique characteristics. The different karyomorphs are a consequence of the presence of Robertsonian rearrangements, such as centric fissions and pericentric inversions, which play an important role in the evolution of Hypostominae. Although variable in relation to the location of constitutive heterochromatin, we observed the presence of banding C in some chromosomes of all karyomorphs, which may indicate the existence of some homology. Another conservative feature is the presence of two pairs of subtelocentric or acrocentric chromosomes carrying 18S rDNA cistrons in the terminal region of the chromosomes. However, we observed the discontinuity of cytogenetic and phylogenetic data, with the formation of different groups (Araras + Indaiatuba and Botucatu + Terra Roxa in cytogenetics, in contrast to Araras + Terra Roxa and Botucatu and Indaiatuba in the phylogeny), suggesting that several derived karyomorphs may be produced from a pluripotent karyomorph as a result of the intrinsic plasticity of the species karyotype. Thus, each new arrangement would be independent in the forms analyzed, as they do not seem to be lineages from the same direct ancestor. Given the above, we believe that the genus *Hypostomus* continues to be one of the most diverse among the Siluriformes, however, we began to understand a little more about the karyotypic diversity of the group by associating different approaches, such as phylogenetic analyses.

## INTRODUCTION

Plecos of the subfamily Hypostominae (Loricariidae) constitute a large group of fish, megadiverse and with complex taxonomies, organized in approximately 45 valid genera and 500 valid species (Fricke et al., 2023). The large variation in characters such as coloration and external morphology (Oyakawa et al. 2005; Zawadzki et al. 2008) and wide distribution in South American rivers means that the genus *Hypostomus* Lacépède 1803 presents the largest number of species already karyotyped. The group of cryptic species known as *Hypostomus ancistroides* is one of the best-represented species in the literature.

In the 15 natural populations of *H. ancistroides* already studied, there are 15 different karyomorphs, usually with 2n = 68 chromosomes (Artoni and Bertollo 1996; Alves et al. 2006; Rubert et al. 2011; Alves et al. 2012; Bueno et al. 2012; Endo et al. 2012; Fernandes et al. 2012; Pansonato-Alves et al. 2013; Traldi et al. 2013). However, different diploid numbers (Maurutto et al. 2012) and even the presence of differentiated systems of sex chromosomes of the type XX/XY (Rocha-Reis et al. 2018), and ZZ/ZW (Lara-Kamei et al. 2017) have been observed. Markers, such as C-banding and the location of ribosomal cistrons, present significant variations, although for fish, especially Siluriformes, the available cytogenetic data is quite limited.

In this study, we present molecular and chromosomal data of four new populations of *Hypostomus ancistroides* and discuss the evolutionary trends observed for this group of cryptic species complex.

## MATERIAL AND METHODS

The material analyzed in this study came from four sampling points within the basins of the rivers Tietê, Mogi-Guaçu, and Grande, all in the state of São Paulo, Southeastern Brazil. We indicate the geographical coordinates of the sampling points in Table 1 and Figure 1.

**Table 1.**
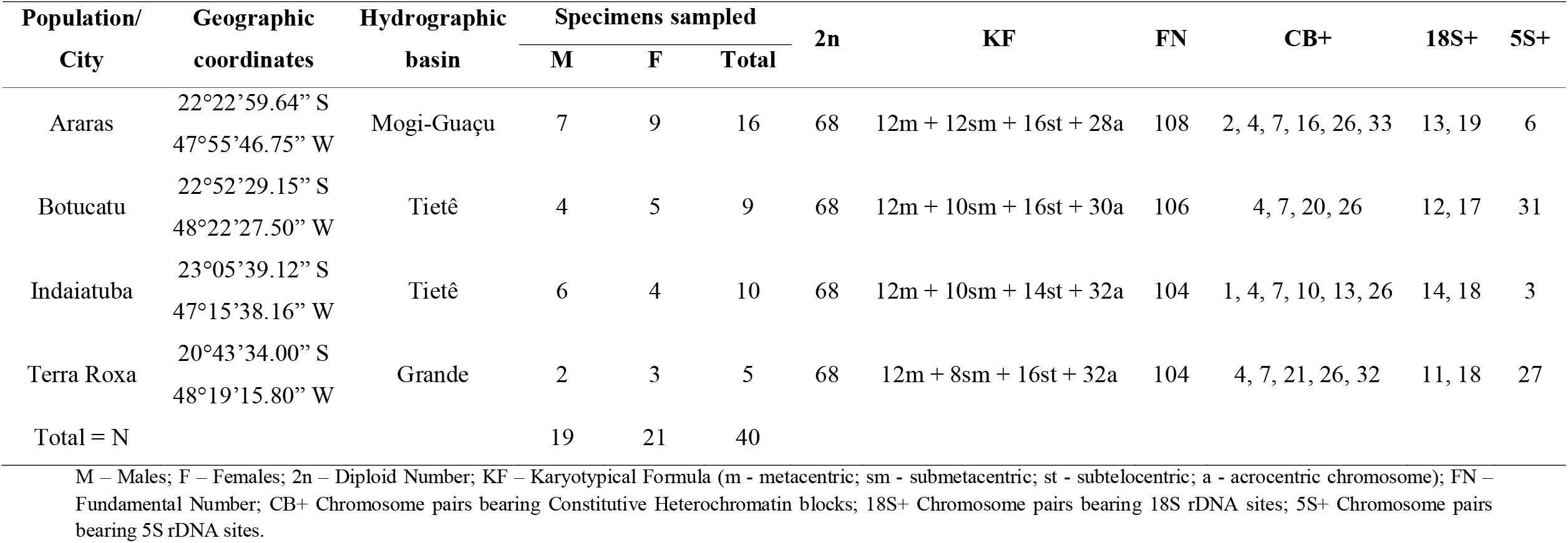
Sampling localities and chromosomal data of the *Hypostomus ancistroides* populations under study.

**Figure 1:**
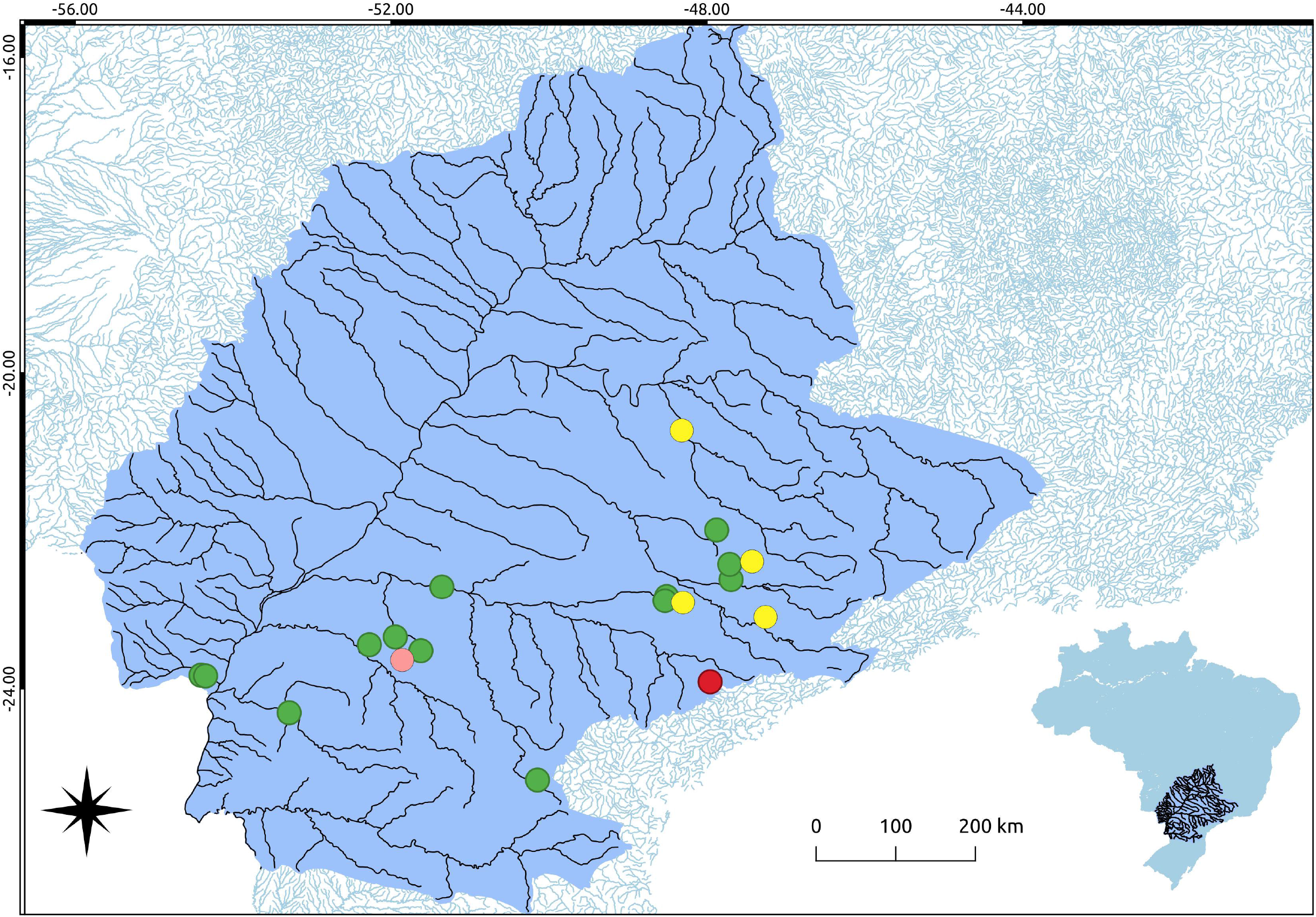
Sampling points along the Tietê, Mogi-Guaçu, Paranapanema and Grande river basins, highlighting in blue the upper Paraná river system in South America. Data from the literature: in green dots, sampled populations of *Hypostomus ancistroides*; in red dot, sampled population of *Hypostomus* aff. *ancistroides* possessing XX/XY sex chromosomal system; in pink dot, sampled population of *Hypostomus* aff. *ancistroides* possessing ZZ/ZW sex chromosomal system; and in yellow dots, populations presented in this paper.

After sampling, we bring the captured specimens alive to the laboratory to euthanize them according to the technical norms of CONCEA (National Council for Control of Animal Experimentation) from Brazil. Taxonomists at the Zoology Museum of the University of São Paulo (MZUSP) had identified all lots as *Hypostomus ancistroides*. Subsequently, we deposited the samples in the Tissue and Cell Suspension Bank and the specimens in the Vertebrate Collection of the Laboratory of Ecological and Evolutionary Genetics of the Federal University of Viçosa, Rio Paranaíba Campus (UFV-CRP), Brazil.

We have obtained the mitotic chromosomes from the renal tissue of individuals by air-drying method (Gold et al. 1990). In addition to the conventional Giemsa staining, we carry out C-banding on *H. ancistroides* chromosomes (Sumner 1972). We performed Fluorescent in situ Hybridization (FISH) (according to Pinkel et al., 1986 and Hamkalo and Elgin, 1991, adapted by Pazza et al., 2006) to localize the sites of rDNA 5S and 18S genes, using as probes the PCR product (*H. ancistroides* DNA was the template). We listed the primers used in this reaction in Table 2. We have labeled the probes with biotin-14-dATP by Nick Translation using the BioNick Labeling System kit (Invitrogen). We have detected the hybridization sites with Cy3 and have mounted the slides with antifade and DAPI (4’-6-diamino-2-phenyl indole). We applied high stringency washes (20% formamide / 0.1xSSC / 15 minutes) and amplified the marker with biotin-conjugated to anti-avidin, incubated in NFDM-PBS (non-fat dry milk) buffer.

**Table 2.**
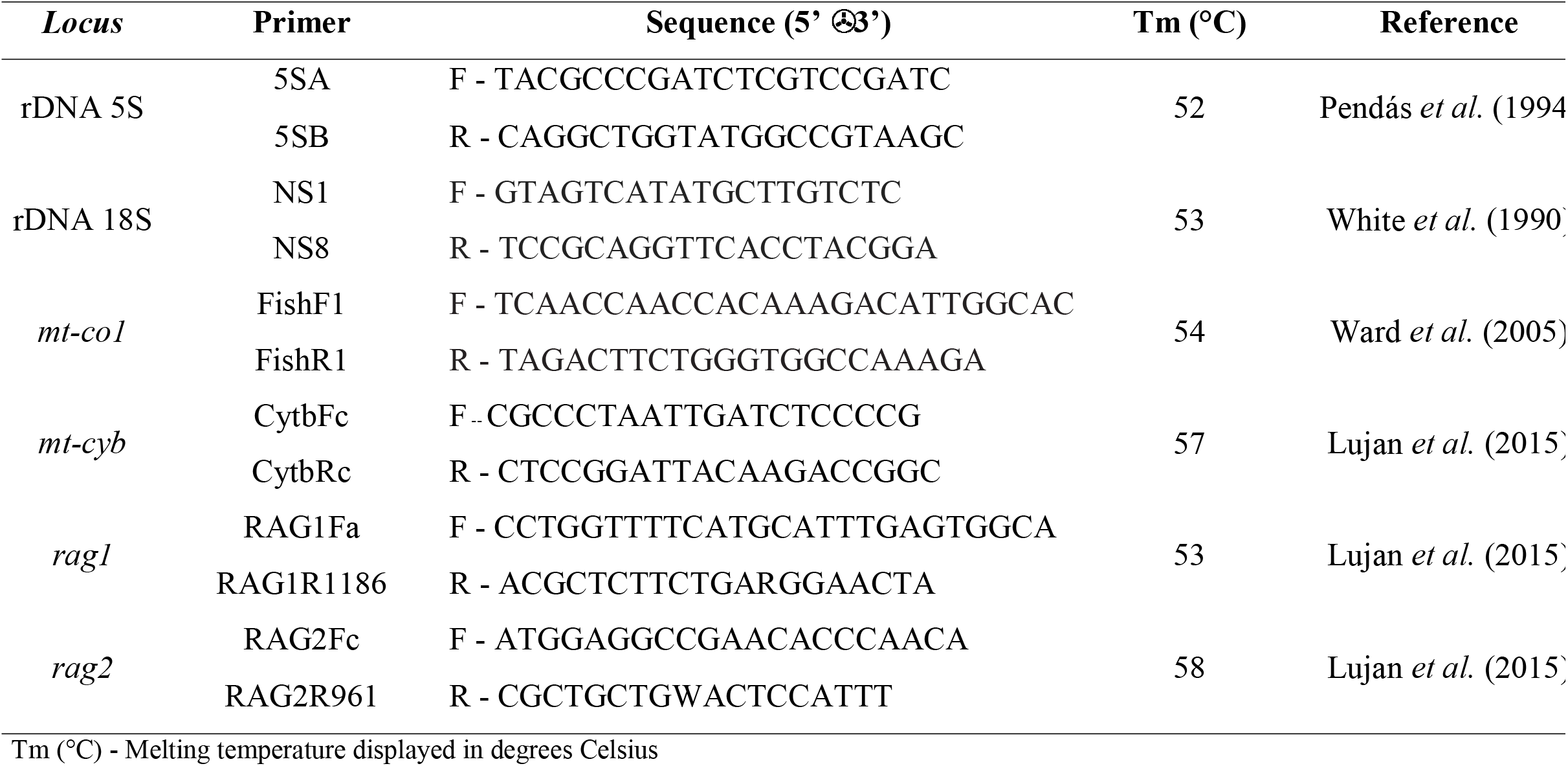
Primers used for amplification of ribossomal, mitochondrial and nuclear genes.

We captured the images with a 3MP definition camera attached to an epifluorescence microscope OLYMPUS BX41. We edited the pictures to assemble the karyotypes using the software GIMP 2.8.14. We classified the chromosomal types by the ratio of the chromosomal arms (Levan et al. 1964).

We sequenced the mitochondrial (Cytochrome c Oxidase I - mt-co1; Cytochrome b - mt-cyb) and nuclear regions (recombination activation gene 1 - rag1; recombination activation gene 2 - rag2) to use in the phylogenetic analysis. We listed the primers in Table 2.

We carried out the Bayesian analysis of the concatenated sequences using the software MrBayes 3.2.6 (Ronquist and Huelsenbeck 2003). We search for the best nucleotide substitution model using PartitionFinder 1.1.1 software for each gene (Lanfear et al., 2012). We evaluated the length of the sampling chain every thousand generations with the Tracer 1.7 software after a 50-million-generation run (Rambaut et al. 2018) to verify the effective sample size (ESS) and strand convergence. We have discarded 25% of the first trees as burn-in with the Tree Annotator software v.1.8. We use FigTree 1.4.2 software to visualize all the phylogenetic trees (Rambaut et al., 2012).

## RESULTS

All specimens have shown 2n=68 chromosomes without supernumerary elements or heteromorphisms related to sex. However, each population possesses a different karyotype with unique characteristics, but traces of homology among them. We summarized all cytogenetic results in Table 1 and illustrated in Figures 2 (karyotypes stained with Giemsa and rDNA 18S FISH), 3 (C-banding), 4 (rDNA 5S FISH), and 5 (population idiograms including all markers obtained).

**Figure 2:**
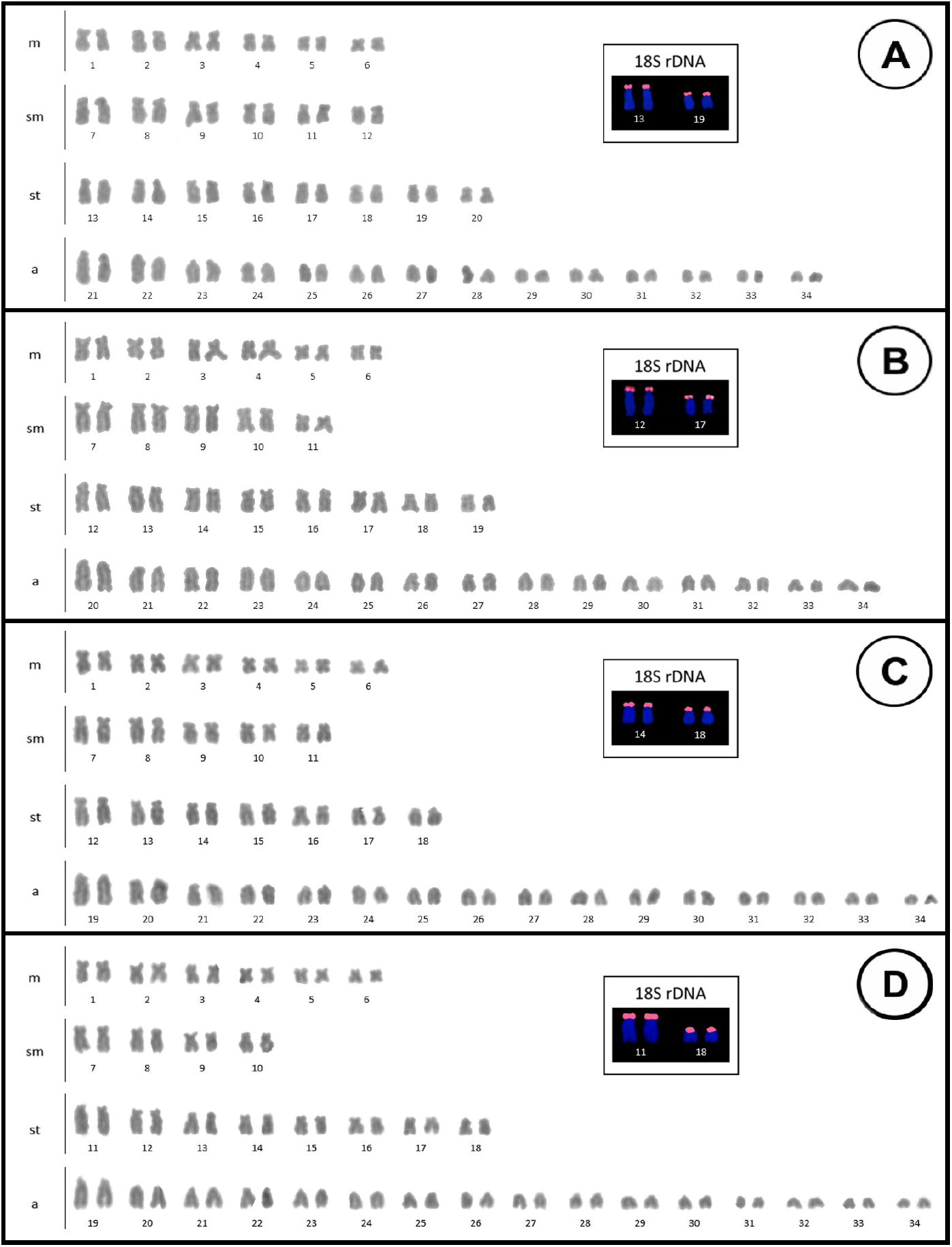
Karyotypes and rDNA 18S gene location (boxes) observed by FISH for the four populations of *Hypostomus ancistroides* presented in this paper: A - Araras, B- Botucatu, C - Indaiatuba, and D - Terra Roxa.

The karyotypes are quite symmetrical and, although different, have a similar conformation in all populations (Figures 2 and 5). In contrast, we have found the 18S rDNA cistrons, always located on subtelocentric chromosomes, in two pairs of similar size. These chromosomal pairs probably correspond to homeologous chromosomes among the different populations (Figures 2 - detail, and 5).

Similarly, we probably observed homeologous pairs with heterochromatic blocks in the four karyomorphs: pairs 4, 7, and 26 (Figures 3 and 5). In these pairs, the heterochromatic blocks may have a different size or be in a different position along the chromosome arm. We also observed other blocks restricted to specific populations, which may constitute populational markers. For example, only in the population from Indaiatuba, we detected C-bands in the first metacentric pair. On the other hand, we have observed a conspicuous C+ block in one of the chromosomes of the acrocentric pair 21, only in the chromosomes of a few individuals from the Terra Roxa population. This feature constitutes a heteromorphism (Figures 3 and 5).

**Figure 3:**
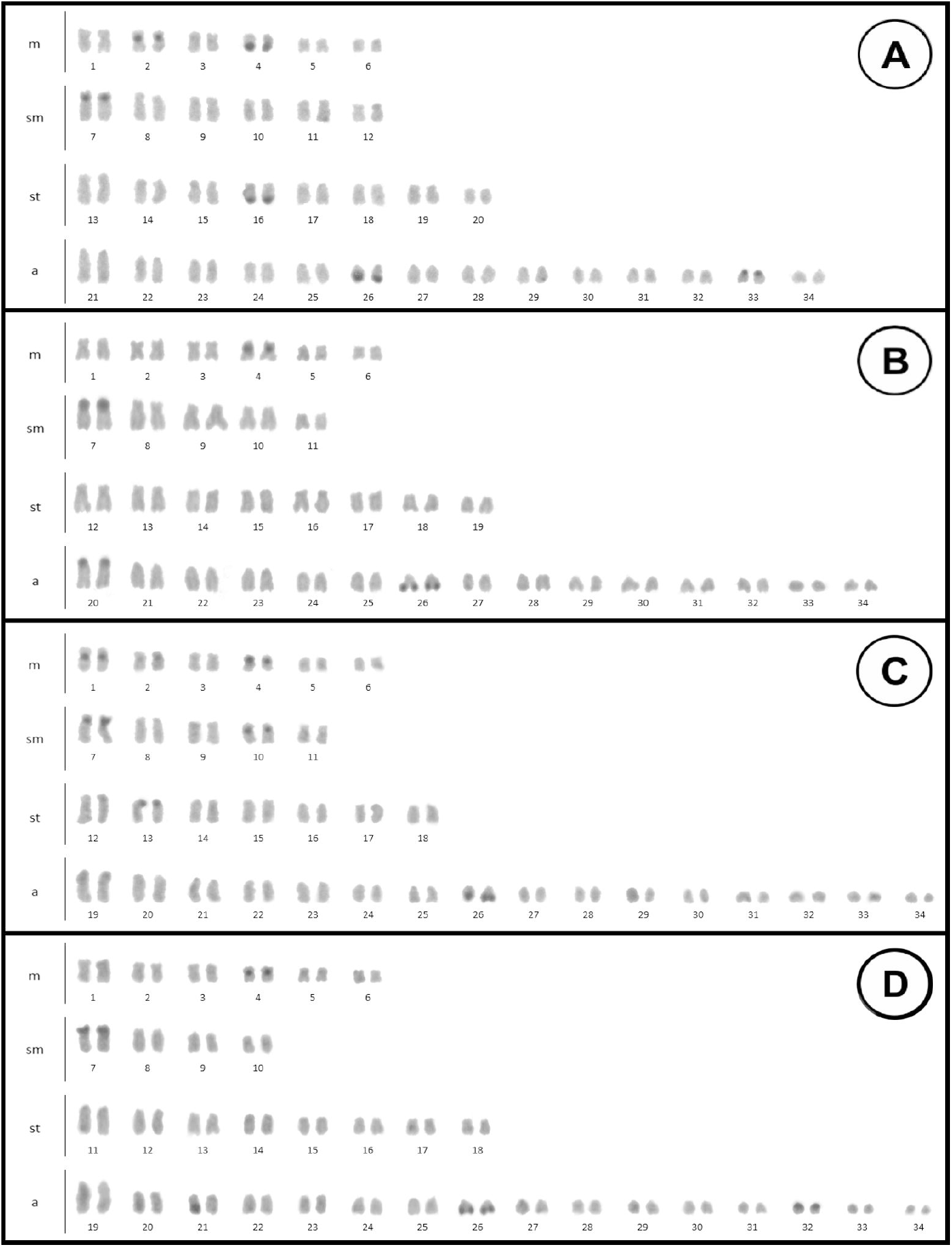
Karyotypes showing the constitutive heterochromatin pattern obtained by C-banding for the four populations of *Hypostomus ancistroides* from: A - Araras, B- Botucatu, C - Indaiatuba, and D - Terra Roxa.

Generally, we observed two phenotypes concerning the location of the rDNA 5S cistrons, although they occur in just one chromosome pair in the four populations studied. The karyomorphs from Araras and Indaiatuba presented the 5S rDNA sites in an interstitial position in the short arm of metacentric chromosomes. The karyomorphs from Botucatu and Terra Roxa presented them in the terminal position on the long arm of the medium acrocentric chromosome pair (Figures 4 and 5).

**Figure 4:**
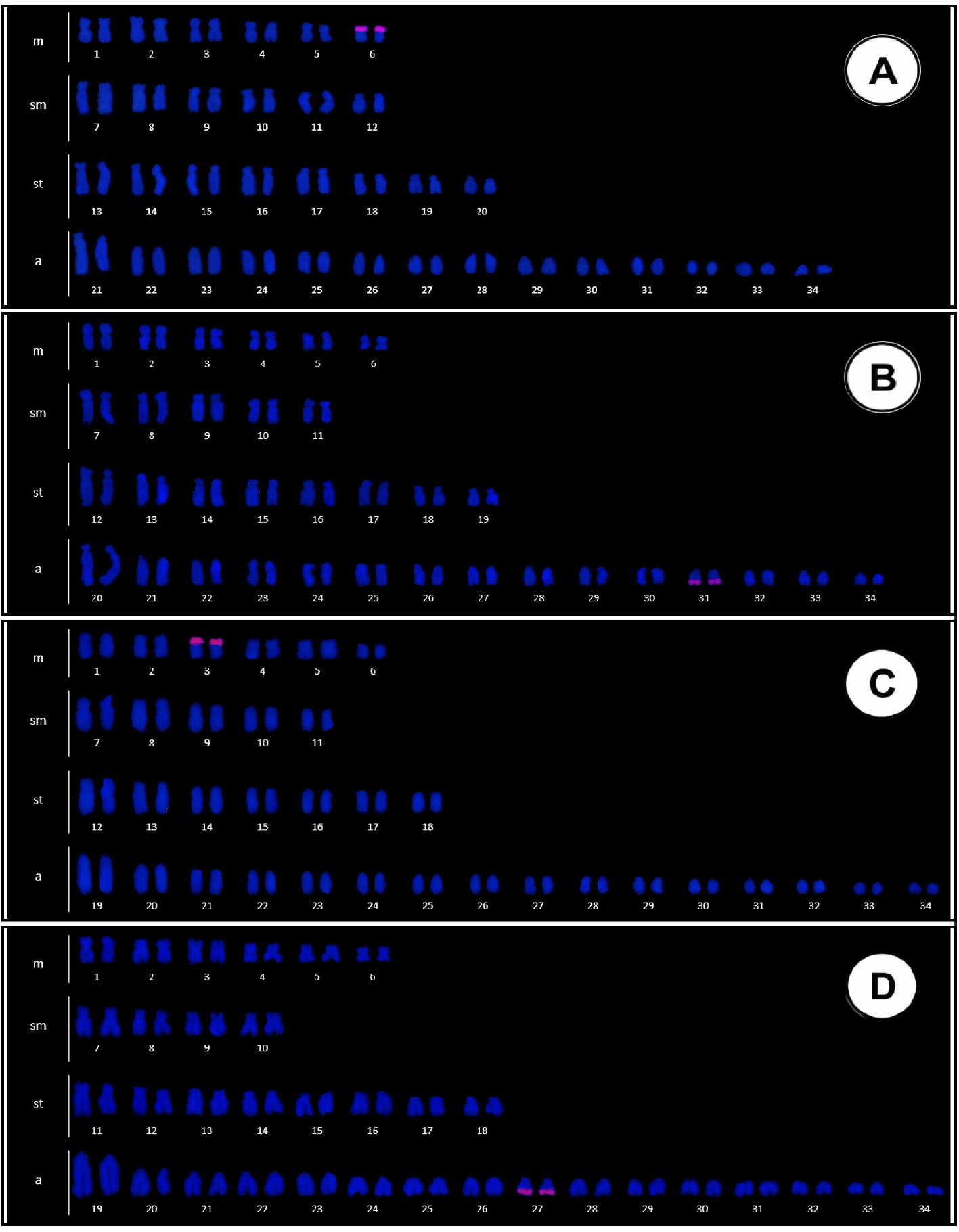
Karyotypes showing the rDNA 5S gene location observed by FISH for the four populations of *Hypostomus ancistroides* presented in this paper: A - Araras, B- Botucatu, C - Indaiatuba, and D - Terra Roxa.

**Figure 5:**
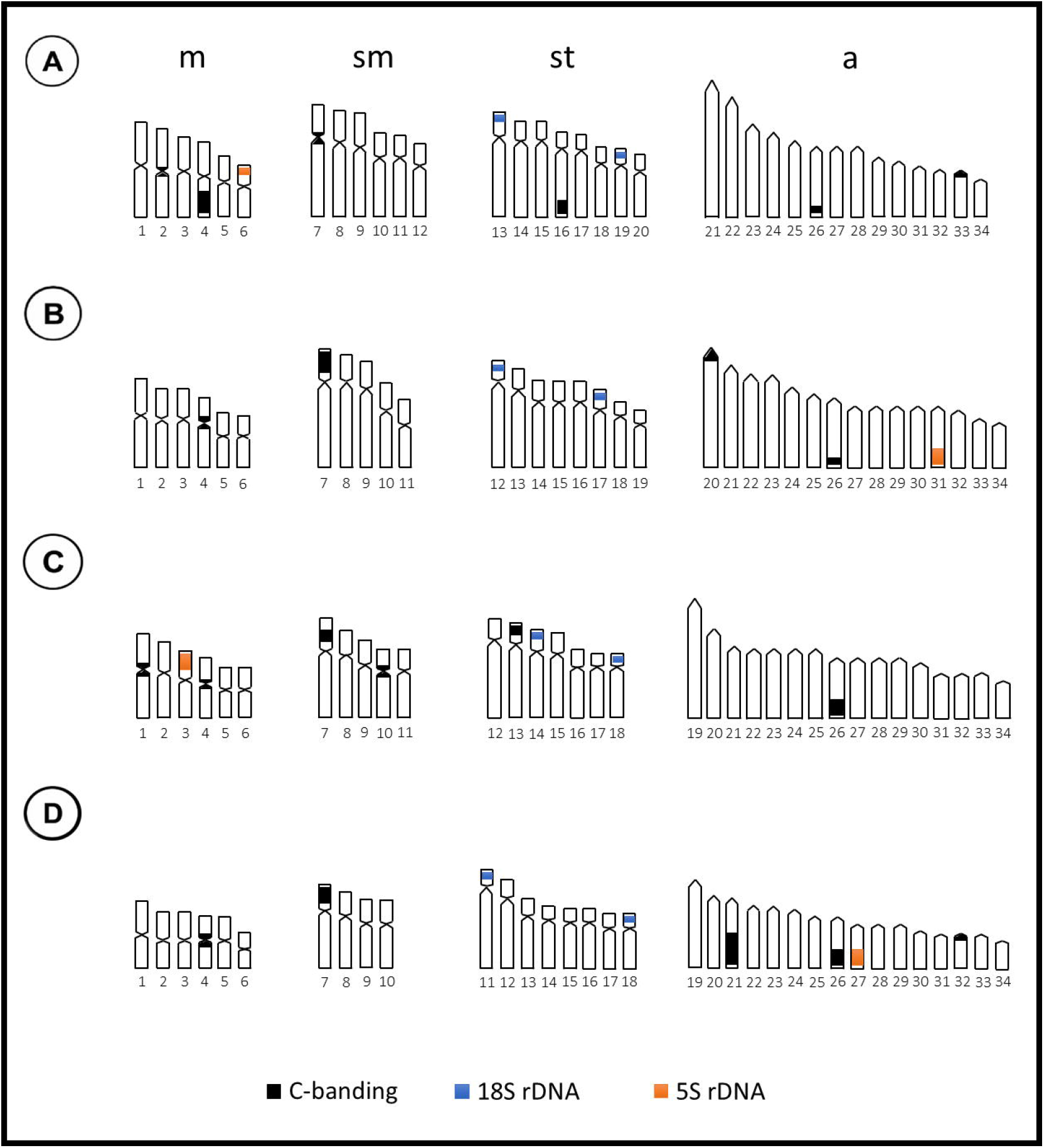
Ideograms summarizing the cytogenetic data observed for the four populations of *Hypostomus ancistroides* presented in this paper: karyotype constitution with karyotypic formula, C-banding, and rDNA 18S and 5S location by FISH. A - Araras, B- Botucatu, C - Indaiatuba, and D - Terra Roxa.

The clusters verified by the localization of the 5S rDNA sites (Figures 4 and 5) are not supported by the molecular data. Bayesian analysis of the concatenated genes (Figure 6), even as other phylogenetic analyses, displays two clusters, one formed by individuals from Araras and Terra Roxa and another by individuals from Botucatu and Indaiatuba. The analyses also show the sharing of haplotypes within each group, but not between the two clusters. The populations from Botucatu and Indaiatuba both share the Tietê River basin. However, Araras and Terra Roxa populations are from several kilometers away and in different river basins, without recent connection.

**Figure 6:**
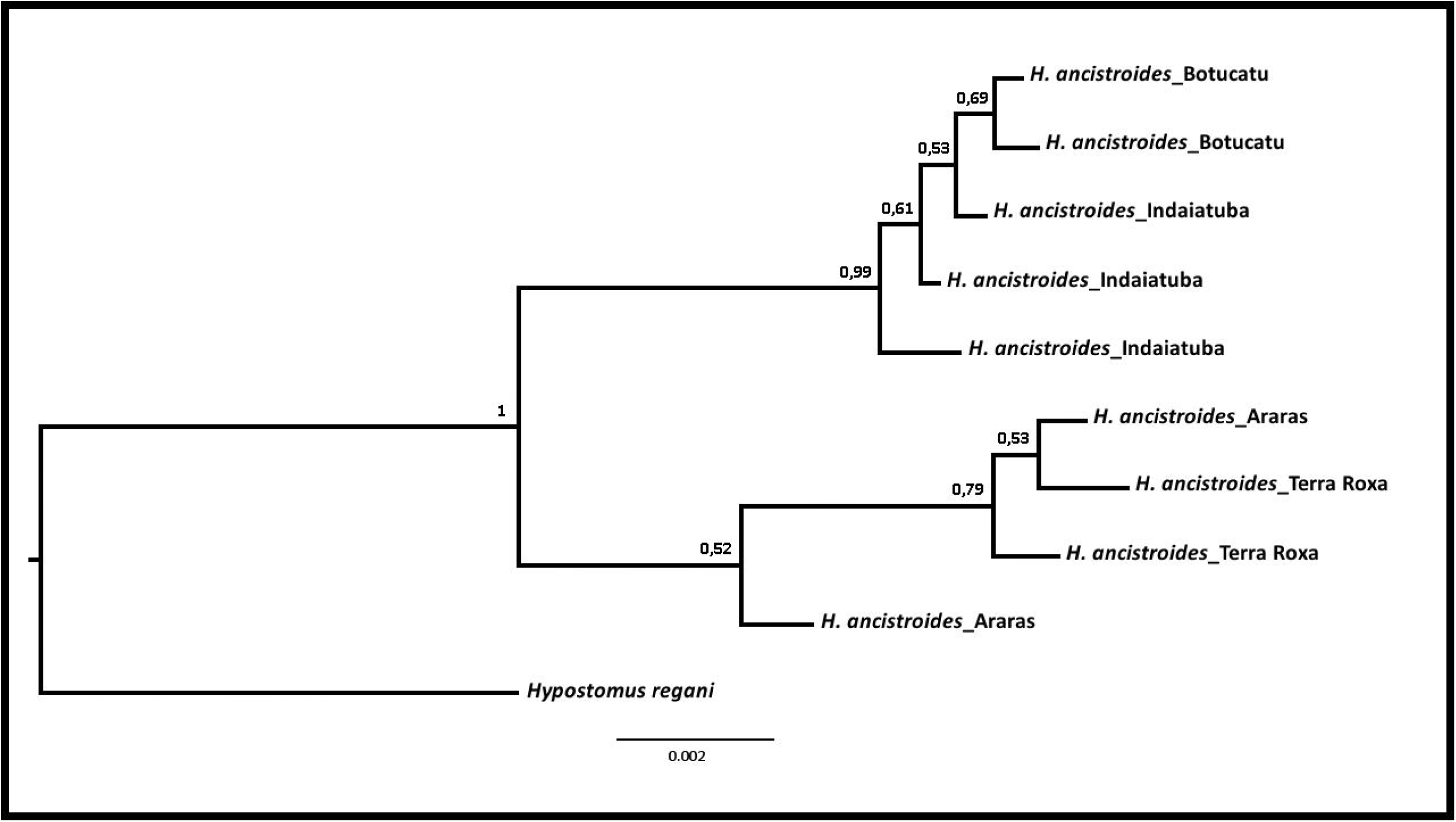
Bayesian tree obtained by concatenation of mitochondrial and nuclear genes sequenced from the four populations of *Hypostomus ancistroides*. Numbers in nodes represent posterior probability values. Sequences from *Hypostomus regani* were used as outgroup.

## DISCUSSION

Britski (1972) reports that *Hypostomus* is the dominant genus of plecos in Brazil, occurring in a multifold variety of freshwater ecosystems (Oyakawa et al., 2005). Despite being the most studied genus from a cytogenetic point of view, the great diversity of this group remains almost unexplored (Rubert et al., 2011) compared to the number of species that are described annually. They exhibit wide karyotypic variation, with species presenting from 64, such as *H. faveolus* and *H. cochliodon* (Bueno et al., 2013, 2014) to 84 chromosomes (*Hypostomus* sp. 2 – Cereali et al., 2008). Artoni and Bertollo (1996) consider this group as not conserved concerning the karyotypic macrostructure.

The maintenance of the diploid number could represent adaptive and ancestral state populations in this study and seems to be a tendency in the group. The four samples we analyzed have shown the same diploid number as the great majority of the studied populations, 2n = 68 chromosomes (Artoni and Bertollo 1996; Alves et al. 2006; Bueno et al. 2012; Rubert et al. 2011; Endo et al. 2012; Alves et al. 2012; Fernandes et al. 2012; Pansonato-Alves et al. 2013; Traldi et al. 2013; Lara-Kamei et al. 2017) (Figures 1 and 4). The exception for this character is an isolated population in the Tibagi River (Maurutto et al. 2012) and a case of a new species of the *H. ancistroides* complex carrying a differentiated sexual chromosomal system, which is quite divergent from the others (Rocha-Reis et al. 2018).

Despite this, we observed differences in the karyotypic formulas (Figures 1 and 4), a consequence of the presence of Robertsonian rearrangements, such as centric fissions and pericentric inversions, which play an important role in the evolution of Hypostominae (Artoni and Bertollo, 2001). The establishment of these rearrangements in populations can be facilitated by the habitat of these animals, as they are often considered organisms that migrate short distances, forming isolated populations that are more conducive to inbreeding (Almeida-Toledo et al., 2000).

The role of chromosomal rearrangements in the diversification of species has been a subject of debate for many years, and there is evidence that unbalanced rearrangements can interfere in gametogenesis, reinforcing the reproductive isolation of karyomorphs by reduction of gene flow, as some species have increased tolerance to chromosomal rearrangements, maintaining polymorphic populations or possessing large karyotype plasticity (Pazza et al. 2018).

Chromosomal variations in *Hypostomus* are not restricted only to the karyotype formulae, occurring also in patterns of location of the constitutive heterochromatin (Figures 3 and 5). Although variable when observing the totality of the karyotype macrostructure, the presence of C-banding in some chromosomes of all karyomorphs may indicate the existence of some homology (Rubert et al. 2011; Fernandes et al. 2012; Maurutto et al. 2012; Pansonato-Alves et al. 2013; Traldi et al. 2013; Lara-Kamei et al. 2017; Rocha-Reis et al. 2018), and may even be considered a phylogenetic sign. In this study, we observed the presence of conserved C+ blocks in pairs of chromosomes 4 (m), 7 (sm) and 26 (a), in a pericentromeric or subterminal region (Figures 2 and 4). Other markings, however, were quite autapomorphic. Individuals of *H. ancistroides* from Terra Roxa displayed polymorphisms related to heterochromatin distribution (Figure 3D - pair 21). The absence of blocks in one of the homologs reveals the likely occurrence of unequal exchanges during cell division, where a part or the entire heterochromatin block is translocated to another chromosome.

Interestingly, such polymorphisms are observed, especially in heterokariotypes, since the major part of the polymorphisms appears in heterozygosity in populations, and, depending on demographic events and evolutionary processes, can be fixed or eliminated over generations, usually after overcoming subdominance (Hoffmann and Rieseberg 2008; Kirkpatrick 2010) or by selection or genetic drift in small populations (Spirito 1998). This indicates not only that there is variation in the population, but it is in an overt process of chromosome evolution.

Another characteristic that seems to be conservative in the group of *Hypostomus ancistroides* cryptic species and could be adaptive is the presence of two subtelocentric or acrocentric chromosome pairs carrying the 18S rDNA cistrons (Figures 2 – details and 5), a trend observed in several studies (see Rubert et al. 2011; Pansonato-Alves et al. 2013; Traldi et al. 2013; Bueno et al. 2014; Lara-Kamei et al. 2017; this study). Furthermore, all these sites are located in the terminal region of the chromosomes (Rocha-Reis et al. 2021).

The dynamics of dispersion of repetitive sequences in chromosomes is often associated with the presence of active transposable elements (TE) in the genomes (Silva-Neto et al. 2015), which could explain the different patterns in the distribution of heterochromatin blocks and rDNAs 5S cistrons in different pairs of chromosomes (Figures 4 and 5), as a plastic characteristic. rDNA sequence analyses have demonstrated the existence of these elements in spacer regions, disseminating gene families in functional copies or pseudogenes (Drouin et al. 1995; Gornung 2013; Rebordinos et al. 2013; Symonová et al. 2013). The dispersion of TEs (and consequently of ribosomal DNA) could then affect the rate of recombination in the genomes and lead to rapid divergence of the karyotype/genome.

The discontinuity between phylogeography constructed from the sequence of four genes (mtDNA and nDNA) and 5S rDNA phenotypes in *H. ancistroides* (Figures 4 and 6) could be a result of convergence in the location of ribosomal cistrons, generated by translocations in both clusters (Araras + Terra Roxa and Botucatu + Indaiatuba). However, the idea that several derived karyomorphs can be produced from one pluripotent karyomorph as a result of the intrinsic karyotype plasticity of the species is more parsimonious, there being multiple possible forms for each type of chromosome character, and the reality that the chromosomes carrying the sites of rDNA 5S are not necessarily homologous to that of the ancestral karyomorph. That is, each new arrangement would be independent in the analyzed forms since they do not appear to be lineages of the same direct ancestor. This would explain not only the distribution pattern of this gene but also the existence of different karyotype formulas and heterochromatic blocks not shared between populations.

The *Hypostomus* genus remains one of the most diverse among the Siluriformes, however, we began to understand a little more about the karyotypic diversity of the group by associating different approaches, such as phylogenetic analyses. It shows us, for example, plastic or homologous characters within karyotypes, and this will certainly help to understand the karyotypic evolution of this specious group.

## Notes

### Competing Interest Statement

The authors have declared no competing interest.

### Summary of Updates

We request that you update the order of authors, title and text of this manuscript. This change is due to the meticulous review carried out by the authors, aiming to make the text more concise and suitable for publication, as suggested by independent reviewers.

